# Recruitment of Neuronal KCNQ2/3 Channels to Membrane Microdomains by Palmitoylation of Alzheimer’s Disease-Related Protein BACE1

**DOI:** 10.1101/2020.09.30.321307

**Authors:** Gucan Dai

## Abstract

β-secretase 1 (β-site amyloid precursor protein (APP)-cleaving enzyme 1, BACE1) plays a crucial role in the amyloidogenesis of Alzheimer’s Disease (AD). BACE1 was also discovered to act like an auxiliary subunit to modulate neuronal KCNQ2/3 channels independent of its proteolytic function. BACE1 is palmitoylated at its carboxyl-terminal region, which brings BACE1 to ordered, cholesterol-rich membrane microdomains (lipid rafts). However, the physiological consequences of this specific localization of BACE1 remain elusive. Using spectral Förster Resonance Energy Transfer (FRET), BACE1 and KCNQ2/3 channels were confirmed to form a signaling complex, a phenomenon that was relatively independent of the palmitoylation of BACE1. Nevertheless, palmitoylation of BACE1 was required for recruitment of KCNQ2/3 channels to lipid-raft domains. Two fluorescent probes designated L10 and S15, were used to label lipid-raft and non-raft domains of the plasma membrane, respectively. Coexpressing BACE1 substantially elevated the FRET between L10 and KCNQ2/3 whereas the BACE1-4C/A quadruple mutation failed to produce this effect. In contrast, BACE1 had no significant effect on the FRET between S15 probes and KCNQ2/3 channels. A reduction of BACE1-dependent FRET between raft-targeting L10 probes and KCNQ2/3 channels by applying cholesterol-extracting reagent methyl-β-cyclodextrin (MβCD), raft-disrupting general anesthetics, or pharmacological inhibitors of palmitoylation all supported our hypothesis of the palmitoylation-dependent and raft-specific localization of KCNQ2/3 channels. Furthermore, mutating the four carboxyl-terminal cysteines (4C/A) of BACE1 abolished the BACE1-dependent increase of FRET between KCNQ2/3 and a lipid raft-specific protein caveolin 1. Collectively, we propose how the AD-related protein BACE1 underlies the localization of a neuronal potassium channel.

## INTRODUCTION

This paper concerns a regulatory interaction of an Alzheimer’s-related enzyme and a potassium (K^+^) channel. Alzheimer’s disease (AD), the most prevalent form of dementia, is characterized by elevated amounts of amyloid plaque in the brain (Perl, 2010; Weller and Budson, 2018). β-secretase 1 (BACE1) is one of the major proteases that cleave the amyloid precursor protein (APP), leading to the accumulation of amyloid β (Aβ) fragments (Vassar et al., 2014). A single mutation in APP that compromises the ability of BACE1 to cleave APP can reduce Aβ generation and slow the progression of AD (Di Fede et al., 2009; Jonsson et al., 2012). Therefore, inhibition of BACE1 has been proposed as a promising therapeutic strategy to treat AD (Vassar and Kandalepas, 2011). Indeed, mice with BACE1 gene knockout, BACE1(-/-), do not exhibit Aβ plaques or develop AD; however, they do show debilitating changes in behavior tests, including spatial memory deficits, and sensorimotor impairments (Cole and Vassar, 2007; Kobayashi et al., 2008). Furthermore, BACE1(-/-) mice exhibit neuronal deficits including axonal organization defects in the hippocampus (Ou-Yang et al., 2018), myelination deficits (Hu et al., 2006) and altered synaptic function and neurotransmitter metabolism (Lombardo et al., 2019). In addition, BACE1(-/-) mice exhibit epileptic seizures, suggesting a possible direct role of BACE1 in regulating neuronal voltage-gated Na^+^ and K^+^ channels (Kim et al., 2007; Lehnert et al., 2016).

Candidate ion channels for regulation by BACE1 include the M-type K^+^ channels, which consist of KCNQ1-5 (Kv7.1-7.5) subunits (Brown and Adams, 1980; Hessler et al., 2015). Two channel genes, *KCNQ2* and *KCNQ3*, are widely expressed in the central nervous system (Wang et al., 1998); they form a heteromeric channel important for transducing sympathetic stimuli, dampening excessive neuronal excitability, reducing excitatory postsynaptic potentials and controlling neuronal firing (Abbott, 2020; Hernandez et al., 2008). KCNQ2/3 channels are abundantly present in the hippocampus and are critical for hippocampal excitability (Carver et al., 2020; Klinger et al., 2011; Shah et al., 2002; Yue and Yaari, 2004). In 2015, BACE1 was reported to interact with KCNQ2/3 channels directly to potentiate M-currents in hippocampus (Hessler et al., 2015). Using patch-clamp electrophysiology, immunoprecipitation, and proximity-ligation assay, BACE1 was found to function as an auxiliary subunit of KCNQ2/3 channels, apparently independent of its proteolytic activity (Hessler et al., 2015). Since KCNQ channel genes are well known for their involvement in several types of epilepsy (Biervert et al., 1998; Singh et al., 1998), this research seemingly provides a novel linkage between KCNQ2/3 channels and the epileptic phenotypes seen in BACE1(-/-) mice. Not surprisingly, it was also found that AD patients are more susceptible to develop epilepsy (Vossel et al., 2017).

The enzymatic activity of BACE1 as an APP protease occurs in membrane lipid raft domains (Cordy et al., 2003; Ehehalt et al., 2003). Lipid rafts are dynamic membrane microdomains containing sphingolipids, cholesterol, and certain proteins (Cornell et al., 2020; Matsubara et al., 2021; Nahmad-Rohen et al., 2020; Pavel et al., 2020; Robinson et al., 2019; Simons and Ikonen, 1997). Rafts are recognized as detergent-resistant membrane fractions, structurally more ordered than non-raft areas; they are signaling hubs orchestrating many cellular events (Lingwood and Simons, 2010; Pike, 2004). Lipid-raft localization of BACE1 biases the processing of APP in the amyloidogenic direction, generating Aβ (Ehehalt et al., 2003; Wang et al., 2021). When APPs are outside of raft domains, they are preferentially cleaved by the non-amyloidogenic α-secretase (Cordy et al., 2006; Kojro et al., 2001; Wang et al., 2021). Similar to other raft-localizing proteins like tyrosine kinases, G protein α-subunits, and purinergic P2X7 receptors, BACE1 uses palmitoylation, in this case, of four cysteines at its carboxyl-terminal end for specific membrane anchoring (Linder and Deschenes, 2007; Vetrivel et al., 2009). However, it is not well understood where the non-enzymatic activities of BACE1 localize, especially for regulating KCNQ channels. These channels do not have known palmitoylation sites, but are reported to interact with palmitoylated scaffold proteins, including A-kinase anchoring proteins (AKAP79 in human) and ankyrin-G (Chung et al., 2006; He et al., 2012; Keith et al., 2012; Zhang and Shapiro, 2016). Furthermore, previous research suggested a spatial overlap in the subcellular localization of BACE1 and KCNQ channels; KCNQ channels are thought to be localized in axons and axon initial segments (AIS) via ankyrin-G, whereas BACE1 is abundant in axons and implicated functionally in the myelination of axons (Buggia-Prevot et al., 2014; Chung et al., 2006; Hu et al., 2006).

To understand the regulation of KCNQ2/3 channels by BACE1, we looked for localization of the BACE1-KCNQ complex in the lipid rafts of cultured cells and provided further evidence supporting the interaction of BACE and KCNQ2/3. We used Förster Resonance Energy Transfer (FRET) to demonstrate the proximity of membrane proteins and measured FRET, in a spectral manner, based on the sensitized emission of FRET acceptor after exciting the FRET donor. Using engineered fluorescent probes derived from short lipidated sequences of protein kinases Lck and Src to label lipid raft and non-raft domains respectively (Chichili and Rodgers, 2007; Myeong et al., 2021; Rodgers, 2002), we showed that KCNQ channels are localized to lipid rafts. This specific localization of KCNQ channels is dependent on the palmitoylation of BACE1. We provide a strategy to determine the localization of proteins in specific lipid microdomains that could be broadly applied to answer similar questions of other membrane protein complexes.

## RESULTS

This article reports the specific localization of neuronal KCNQ2/3 channels in membrane microdomains (lipid rafts), controlled by formation of a complex with BACE1.

### FRET between KCNQ2/3 and BACE1 consistent with their close assembly

BACE1 interacts with KCNQ2/3 channels directly, demonstrated by previous research using protein co-immunoprecipitation and other biochemical methods (Hessler et al., 2015). Since some KCNQ channels can assemble with single-transmembrane auxiliary subunits e.g., KCNE for cardiac KCNQ1 channels (Abbott, 2020; Sun and MacKinnon, 2020); BACE1, which is also a single-transmembrane protein, has been considered by analogy as a novel auxiliary subunit for KCNQ2/3 channels (Hessler et al., 2015). Atomic details of the interaction between BACE1 and KCNQ2/3 are largely unknown. Here, we further tested this interaction using FRET between a fluorescent noncanonical-amino acid L-Anap (Chatterjee et al., 2013; Kalstrup and Blunck, 2013) (Fig. S1) incorporated into the carboxyl-terminus of a mouse BACE1 and a yellow fluorescent protein (YFP) fused to the carboxyl-terminus of both human KCNQ2 and KCNQ3 subunits all expressed in a cell line. KCNQ2 and KCNQ3 subunits form heteromeric channel complexes and were expressed together throughout this paper in the tsA201 cell line derived from human embryonic kidney cells.

BACE1 consists of an extracellular (amino-terminal) proteolytic domain, one helical transmembrane domain and a short intracellular (carboxyl-terminal) region (Fig. 1A). Four cysteines, one near the cytoplasmic end of the transmembrane domain(C474) and three within the intracellular region (C478, C482, C485) are S-palmitoylated, conferring the lipid raft-localization of BACE1. Mutating these cysteines is reported to reduce the raft localization of BACE1 and amyloid burden but does not abolish the processing of APP by BACE1 (Andrew et al., 2017; Vetrivel et al., 2009). The DXXLL motif in the carboxyl-terminus of BACE1 (Fig. 1A) is involved in the endocytosis and retrograde transport of BACE1 to the trans-Golgi network (Pastorino et al., 2002; Toh et al., 2018). Mutating this motif increases the plasma membrane localization of BACE1 (Pastorino et al., 2002). In addition, the transmembrane and carboxyl-terminal of BACE1 are highly conserved (almost identical) among different species.

**Figure 1.**
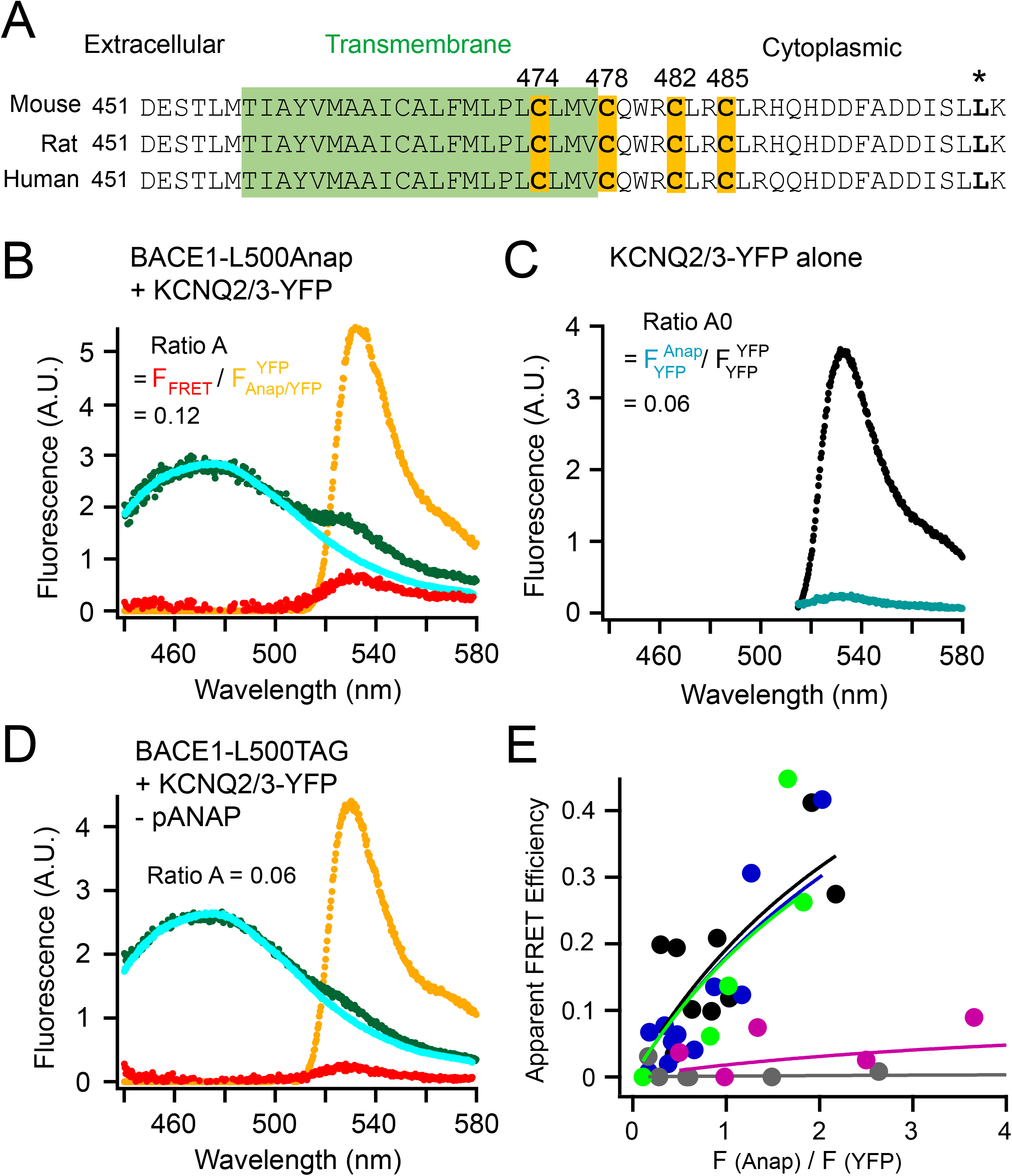
FRET between L-Anap incorporated in BACE1 and YFP-tagged KCNQ2/3 channels. **(A)** Sequence alignment of mouse, rat, and human BACE1 proteins highlighting the four cysteines for palmitoylation and the site of leucine 500 (*). The transmembrane region is in green. **(B)** Representative emission spectra used to calculate FRET between BACE1-L500Anap and KCNQ2/3-YFP. The cyan (donor only) and green spectra (both donor and acceptor) are generated by the Anap excitation wavelength; the yellow spectrum (with both donor and acceptor) by YFP excitation; the red spectrum is the subtraction of the cyan spectrum from the green spectrum. The ratio of the peak intensity of the red spectrum versus that of the yellow spectrum (Ratio A = F_FRET_ / F_Anap/YFP_ ^YFP^) after subtracting the crosstalk (Ratio A0 = F_YFP_ ^Anap^/ F_YFP_ ^YFP^ shown in the panel C) is proportional to the amount of FRET detected. **(C)** Representative emission spectra of KCNQ2/3-YFP-only cells using direct YFP excitation (large spectrum, F_YFP_ ^YFP^) and using excitation for Anap (small spectrum, F_YFP_ ^Anap^) for calculating the Ratio A0. **(D)** Representative emission spectra showing the negative control condition when the BACE1-L500TAG and KCNQ2/3-YFP were transfected but in the absence of the pANAP plasmid. **(E)** Relationship between the apparent FRET efficiency and the ratio of the peak intensity (at around 475 nm) of the green spectrum divided by that of the yellow spectrum (at 530 nm) as in panel B or D in the following conditions shown in different colors: no pANAP (grey), the following conditions all with pANAP; wild-type BACE1 (black), mutant BACE1-4C/A (blue), with added 2-BP (green), and with KCNH instead of KCNQ2/3 (magenta). Each point represents data from one cell.

Considering the small size of the intracellular carboxyl-terminal part of BACE1, fusing a large GFP-derived fluorescent protein might perturb the natural properties of BACE1, so a fluorescent noncanonical amino acid L-Anap was incorporated at the carboxyl-terminal end of the BACE1. An amber stop-codon (TAG) suppression strategy (Chatterjee et al., 2013) was used to incorporate L-Anap at the leucine 500 (L500) position within the DXXLL motif of BACE1 (asterisk, Fig S1). To express BACE1-L500Anap in tsA201 cells, cells were transiently co-transfected with the BACE1-L500TAG cDNA and a plasmid pANAP encoding the orthogonal amber suppressor tRNA/aminoacyl-tRNA synthetase (aaRS) pair for L-Anap and incubated overnight with culture medium supplemented with 20 μM of the membrane-permeable methyl-ester form of L-Anap (Gordon et al., 2018; Zagotta et al., 2016). When the translation machinery arrives at the TAG stop-codon site, translation continues by including L-Anap and, avoiding mRNA chain termination. This strategy has previously been shown to achieve satisfactory incorporation of L-Anap to exogenously-expressed recombinant membrane proteins (Dai et al., 2019; Dai et al., 2021; Dai and Zagotta, 2017). In our experiments, cells grown without applying pANAP plasmid (tRNA/aaRS), but with BACE1-L500TAG cDNA transfected and L-Anap in media were used as negative controls for the FRET experiments.

The Anap emission spectrum overlaps well with the excitation spectrum of YFP, making them a practical FRET pair (Chatterjee et al., 2013; Dai et al., 2019) with an estimated Förster radius R_0_ of 47 Å. A high FRET would indicate close proximity of the two proteins. FRET between L-Anap and YFP was determined by calculations from the emission spectra of both fluorophores measured using a spectrometer attached to the microscope (see Methods). This type of spectrum-based FRET directly measures the degree of the sensitized emission of the FRET acceptor after exciting the FRET donor. The sensitized-emission method also includes corrections for the bleed-through and the crosstalk due to spectral properties of FRET donors and acceptors. Cells that expressed BACE1-L500TAG alone were used to generate a template L-Anap emission spectrum (spectrum in cyan, Fig. 1B) to correct for the bleed-through of the Anap emission in the wavelength range for YFP emission. In addition, cells that expressed only KCNQ2/3-YFP channels (FRET acceptor alone) were used to determine a crosstalk efficiency for FRET: the ratio of peak intensities of a YFP emission spectrum by the Anap excitation versus the YFP emission directly excited by the YFP excitation. This ratio called Ratio A0 was a constant, which equaled 6% for the Anap/YFP pair (Fig. 1C). The 6% crosstalk efficiency had to be subtracted from the FRET spectrum. To determine the FRET spectrum, the fluorescence spectrum was measured when both labeled BACE1 and KCNQ2/3-YFP channels were present (spectrum in green, Fig. 1B). The cyan Anap template spectrum in Fig. 1B was then scaled and subtracted from the green spectrum, yielding the resulting F_FRET_ spectrum (spectrum in red). This subtraction was to correct for the abovementioned bleed-through. Furthermore, the peak intensity of the F_FRET_ spectrum was then divided by the peak intensity of the YFP emission spectrum directly excited by the YFP excitation (spectrum in yellow), generating the ratio called Ratio A. This noticeable difference between the values of the Ratio A (Fig. 1B) versus the Ratio A0 (Fig. 1C) was proportional to the apparent FRET efficiency detected, and suggested that a population of Anap-labeled BACE1 molecules was in close proximity to KCNQ2/3-YFP channels. Finally, as a negative-control experiment, fluorescence spectrum was measured when both the BACE1-L500TAG and KCNQ2/3-YFP channels were co-transfected, but in the absence of the pANAP plasmid (Fig. 1D). In this case, the Ratio A had a similar value as the Ratio A0, suggesting that the nonspecific FRET level was negligible between the free L-Anap in the cell and the KCNQ2/3-YFP.

The calculated apparent FRET efficiency (E_app._) varied with the relative amount of L-Anap-incorporated BACE1 relative to the KCNQ2/3-YFP expressed in each cell (see Methods). The E_app._ correlated positively with the ratio of Anap versus YFP peak fluorescence intensity, and in theory would be saturable to a true FRET efficiency (E_max._) at high values of this fluorescence ratio (Takanishi et al., 2006) (Fig. 1E). Such a positive correlation is typical for ensemble FRET with variable fluorophore stoichiometries (Takanishi et al., 2006; Zacharias et al., 2002). With a low expression of the FRET donor BACE1-L500Anap compared to the YFP, this relationship looked more linear. Note that the fluorescence ratio F(Anap) / F(YFP) here in Fig 1E is not equal to the molar ratio of Anap versus YFP, but rather is also dependent on our optical apparatus and on the FRET efficiency, since FRET decreases Anap fluorescence intensity while increasing that of YFP. Essentially, there were different populations of fluorescent molecules regarding FRET: donors alone (D), acceptors alone (A), and donors paired with acceptors (DA). Since detailed information about stoichiometry, contact site and affinity of the BACE1 and KCNQ2/3 interaction was lacking, fitting the relationship between the F(Anap) / F(YFP) ratio and the apparent FRET efficiency using published models with model-dependent assumptions would be unsuitable or oversimplified (Bykova et al., 2006; Cheng et al., 2007). Therefore, a nonlinear saturable isotherm was used for an empirical fitting of the data (Zacharias et al., 2002) (see Methods). Using this fitting evaluated by chi-square tests (chi-square < 0.1 in all cases), the E_max._ was estimated with a value of 0.88 between BACE1-L500Anap and KCNQ2/3-YFP, indicating a upper limit of the apparent FRET efficiency (Fig. 1E). This E_max._ predicts a distance of 34 Å based on the Förster equation, consistent with a close assembly. Later in the paper, we also used this spectral FRET analysis for the cyan fluorescent protein (CFP) and YFP pairs.

The FRET efficiency between BACE1-L500Anap and KCNQ2/3-YFP did not change apparently when the four palmitoylatable cysteines in BACE1 were mutated to alanines (4C/A) or when cells were incubated with the palmitoylation inhibitor 2-bromopalmitate (Davda et al., 2013) (0.1 mM 2-BP) (see conditions indicated by different colors in Fig. 1E and its legend). Negative-control cells lacking transfected pANAP plasmid showed negligible FRET (E_max._ = 0.006) even though visible Anap fluorescence was present due to nonspecific entry of L-Anap to cells. Furthermore, another voltage-gated potassium channel KCNH (a zebrafish ELK) (Dai and Zagotta, 2017), that was considered to be structurally similar to KCNQ but have no reported interaction with BACE1, showed a smaller FRET (E_max._ = 0.1). These results, together with previously-published work (Hessler et al., 2015), suggested that BACE1 and KCNQ2/3 channels form a protein complex that is relatively independent of the palmitoylation of BACE1.

### BACE1-dependent proximity between KCNQ2/3 and the raft-protein caveolin1

If KCNQ2/3 forms a complex with BACE1, is it localized to lipid-rafts? We used the protein caveolin 1, a principal component of caveolae localized to lipid rafts (Li et al., 1996; Williams and Lisanti, 2004; Zacharias et al., 2002), to answer this question. Caveolin 1 often dimerizes at its amino-terminal region and uses the palmitoylation of three cysteines at its carboxyl-terminus to target to lipid rafts (Williams and Lisanti, 2004; Zacharias et al., 2002). A carboxyl-terminal CFP-tagged caveolin 1 was coexpressed with KCNQ2/3-YFP channels in tsA cells, and their FRET was measured using spectral FRET. If BACE1 could recruit KCNQ2/3 channels closer to the lipid-raft microdomains, a higher FRET between KCNQ2/3-YFP and caveolin 1-CFP would be expected reasonably (Fig. 2A). Again, using the same strategy as abovementioned, the Ratio A0 for the CFP/YFP pair was determined to be 0.136 (Fig. 2E), and the apparent FRET efficiency increased as the ratio of CFP/YFP fluorescence increased (Fig. 2F). The relationship between the apparent FRET efficiency and the CFP/YFP ratio was also fitted empirically with the saturable nonlinear isotherm (Fig. 2F) to estimate the upper limit of FRET efficiency. Without BACE1, the FRET between caveolin 1-CFP and KCNQ2/3-YFP was noticeable (E_max._ = 0.13, Fig 2B, F). Lacking additional evidence of direct complex formation, it is possible that caveolin 1 and KCNQ2/3 might not interact directly like BACE1 with KCNQ2/3. Instead, they might FRET nonspecifically because of a sufficiently high local density (Takanishi et al., 2006; Zacharias et al., 2002). However, with the coexpression of BACE1, there was a large increase in the FRET intensity (E_max._ = 0.25, Fig. 2C, F). In contrast, with coexpression of the BACE1-4C/A mutant construct, there was no increase in FRET, and the relationship between the apparent FRET efficiency and the donor/acceptor intensity ratio superimposed on that seen without BACE1 (E_max._ = 0.11, Fig. 2D, F). Comparison of these conditions was best done at a fixed ratio of F(CFP) / F(YFP), so the representative FRET spectra chosen in Fig. 2 B-D had a similar ratio of CFP/YFP peak fluorescence intensity. Together, these results suggested that, with the help of BACE1, overexpressed KCNQ2/3 channels were drawn to lipid-raft domains where they could FRET more efficiently with raft-specific caveolin 1. Endogenous BACE1 is reported to be present in human embryonic kidney cells (based on the Human Protein Atlas: proteinatlas.org), but in our overexpression system, there may have been too little endogenous BACE1 or other scaffold proteins to occlude the actions of additional overexpressed BACE1.

**Figure 2.**
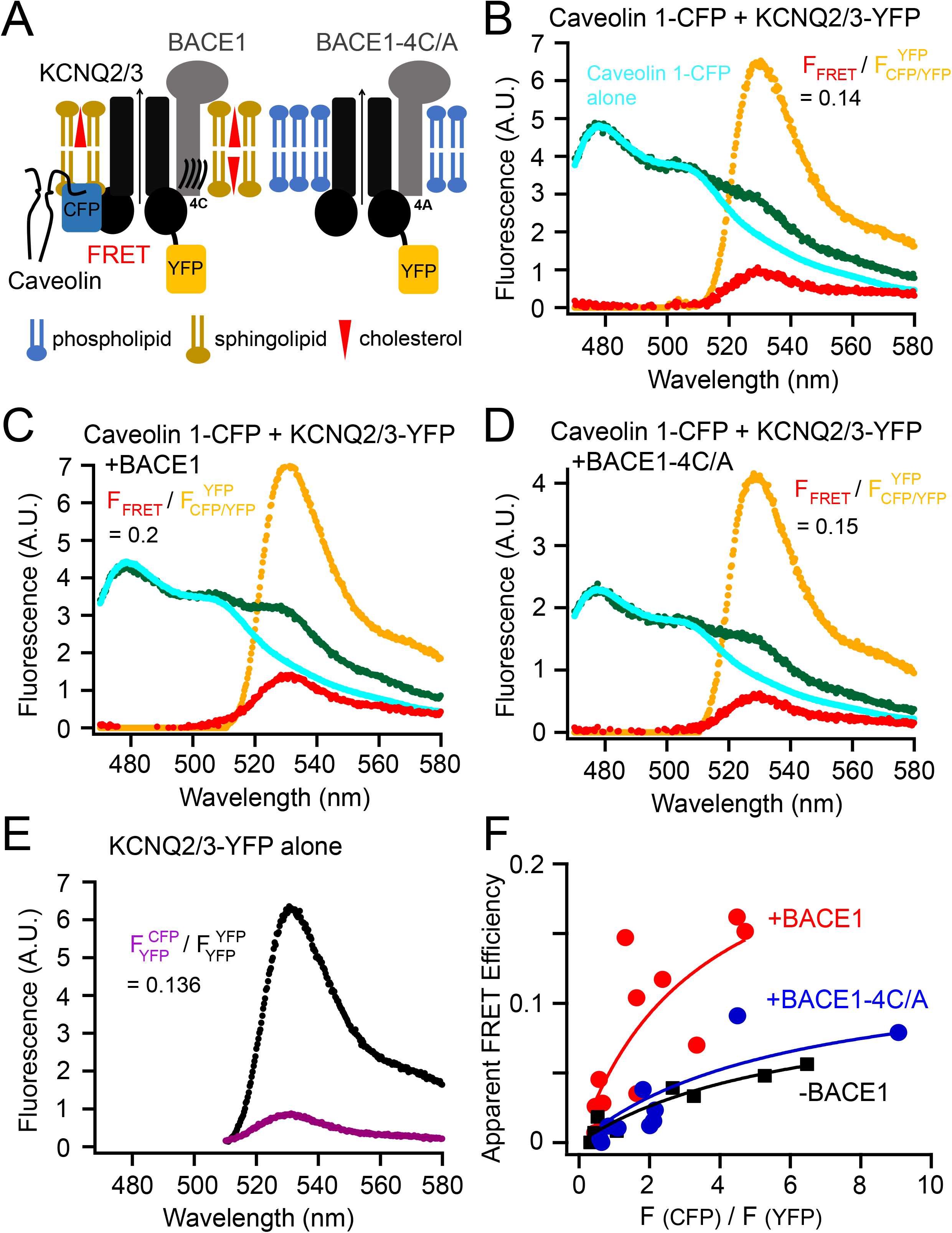
FRET between Caveolin 1-CFP and KCNQ2/3-YFP channels. **(A)** Cartoon illustrating the probing of the localization of KCNQ2/3 channels using the lipid raft-specific protein caveolin 1. **(B-D)** Representative emission spectra for calculating FRET between caveolin 1-CFP and KCNQ2/3-YFP. The cyan and green spectra are generated by illumination of the CFP excitation wavelength; the yellow spectrum by YFP excitation; the red spectrum is the subtraction of the cyan spectrum from the green spectrum. The equation with the numbers shown indicate the Ratio A. **(E)** Representative emission spectrum of KCNQ2/3-YFP alone cells using direct YFP excitation versus the spectrum (smaller amplitude) from the same sample by the excitation for CFP. The number following the equation is the Ratio A0. **(F)** Relationship between the apparent FRET efficiency and the ratio of peak intensities of CFP versus YFP for the FRET between KCNQ2/3 and caveolin 1 in the conditions with or without BACE1 or with BACE1-4C/A.

### Using fluorescent probes that target lipid raft and non-raft domains

To further study the lipid raft-localization of KCNQ2/3 channels, we turned to two fluorescent probes that can be targeted to liquid-ordered and -disordered domains at the plasma membrane (Fig. 3A). The L10 probe derives from the first 10 amino acids of the amino terminus of Lck kinase, which contains two palmitoylation sites for localizing to the ordered lipid-raft domains (Chichili and Rodgers, 2007; Myeong et al., 2021; Rodgers, 2002). In contrast, the S15 probe derives from the first 15 amino acids from the amino terminus of Src kinase, which contains one myristoylation site, for localizing to the disordered non-raft domains (Chichili and Rodgers, 2007; Myeong et al., 2021; Rodgers, 2002).

**Figure 3.**
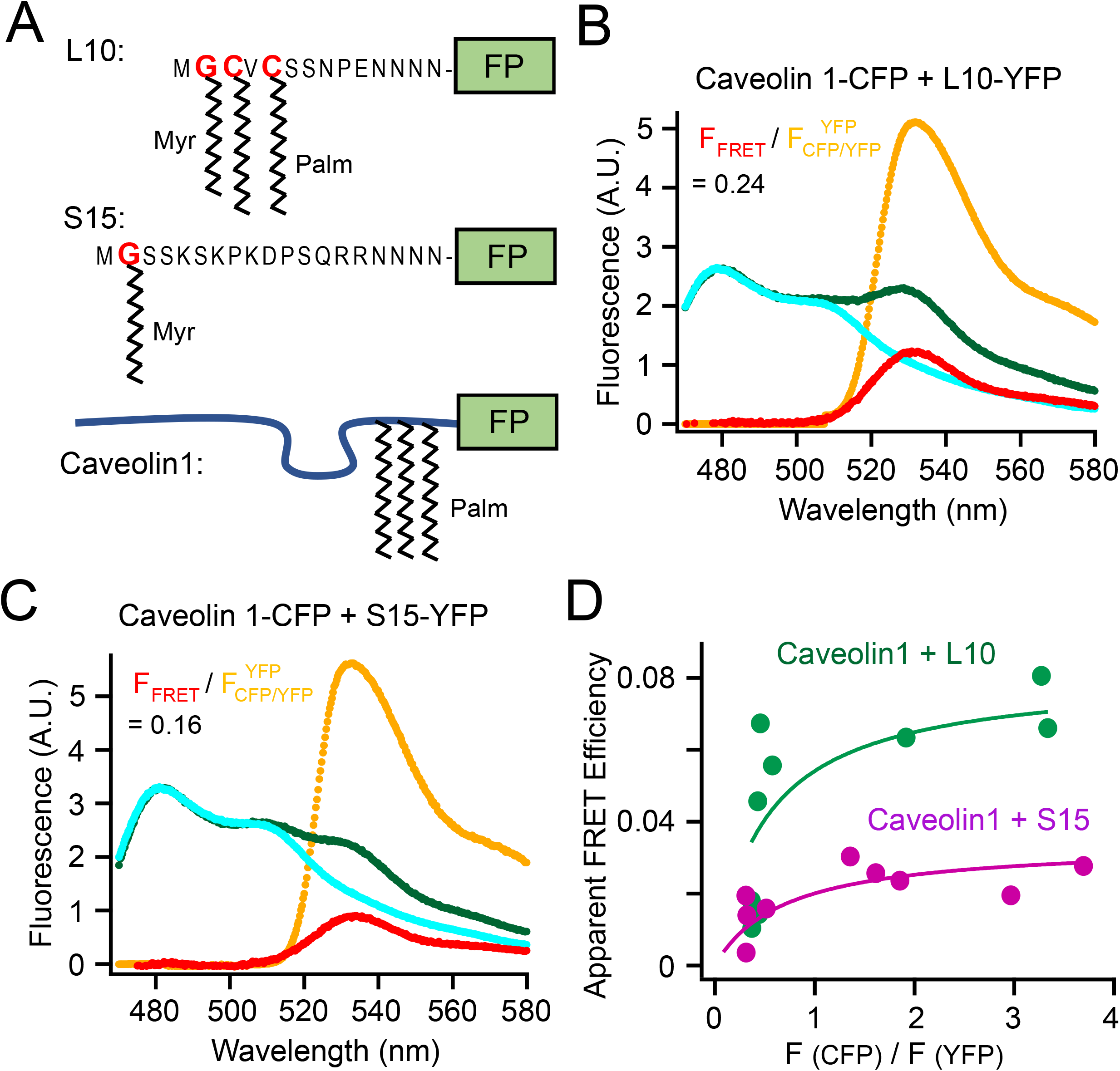
Fluorescent probes that target to lipid rafts and non-rafts. **(A**) Schematic illustration of L10, S15 as well as caveolin 1. Sites of myristoylation and palmitoylation are shown. The Poly-N sequence are the flexible linker connecting to a fluorescent protein (FP), CFP or YFP in this paper. **(B)** Representative emission spectra for the FRET between caveolin 1-CFP and L10-YFP. **(C)** Representative emission spectra for the FRET between caveolin 1-CFP and S15-YFP. **(D)** Relationship between the apparent FRET efficiency and the ratio of peak intensity of CFP versus YFP for the FRET pairs shown in panels B and C.

Validation experiments were first performed comparing the FRET between the Caveolin 1-CFP and L10-YFP pair with the FRET between the Caveolin 1-YFP and S15-YFP FRET pair (Fig 3 B-D). As expected, FRET between L10 and Caveolin 1 was much higher than that between S15 and Caveolin 1, consistent with that the L10 probes localize to raft microdomains where Caveolin 1 is abundant, while S15 localize to non-raft areas (Myeong et al., 2021; Zacharias et al., 2002). Plotting the relationship between the apparent FRET efficiency versus the CFP/YFP ratio suggested a noticeable saturation curve, which might indicate a clustering of fluorescent molecules in a small area (Zacharias et al., 2002). Since lipid rafts are considered small (< 250 nm in diameter), conventional fluorescence microscopy that is limited by the diffraction limit of light cannot visualize them directly. Our FRET method provides an alternative to measure the colocalization of proteins in small raft areas in a relatively high-throughput manner.

Next, L10 and S15 probes were used to test for possible colocalization with KCNQ2/3 channels and to learn how it might be regulated by BACE1. L10-CFP probes were coexpressed with KCNQ2/3-YFP channels without BACE1. The FRET between L10-CFP and KCNQ2/3-YFP was small (E_max._ = 0.04, black curve, Fig 4 A). Coexpression of BACE1 significantly elevated the FRET (E_max._ = 0.27, red curve), whereas coexpression of the mutant BACE1-4C/A did not (E_max._ = 0.06, blue curve). However, using the same experimental settings, neither BACE1 nor the mutant BACE1-4C/A construct had an effect on the FRET between S15-CFP and KCNQ2/3-YFP (Fig 4 B). In addition, a higher FRET efficiency for KCNQ2/3-YFP with S15-CFP (E_max._ = 0.14) than with L10-CFP (E_max._ = 0.04) without the coexpression of BACE1, suggests that KCNQ2/3-YFP channels by themselves were more distant from L10 than from S15. In other words, in the absence of BACE1, KCNQ2/3 might be distributed more in the non-raft areas at least in our overexpression system, possibly due to their lack of known palmitoylation sites. Nevertheless, other fluorescence methods such as super-resolution microscopy would help to further test this idea. In addition, our conclusion is predicated on the assumptions that CFP fluorophores in L10 and S15 probes have comparable distances to the membrane and dipole orientations. To summarize, overexpressed KCNQ2/3 channels initially showed somewhat greater proximity with S15 than with L10 considering that endogenous BACE1 levels are low, however, overexpressing BACE1 apparently increased the localization of KCNQ2/3 channels to lipid rafts. With BACE1, KCNQ2/3 showed higher FRET with L10 than with S15 presumably due to a direct interaction between KCNQ and BACE1. Mutating the 4 cysteines in BACE1 to alanine eliminated this effect, corroborating a critical role of palmitoylation for the lipid raft-localization of both BACE1 and its partner KCNQ2/3.

**Figure 4.**
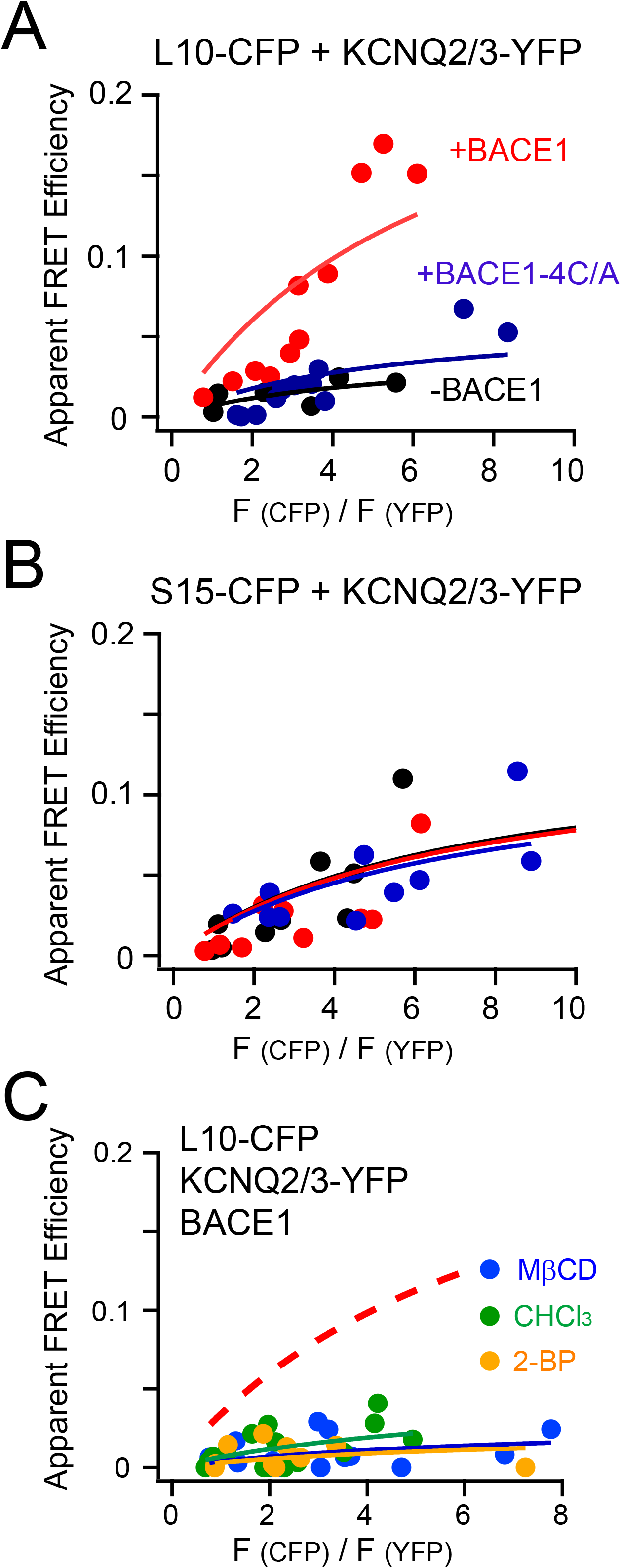
Using L10, S15 to probe the localization of KCNQ2/3 channels. Relationship between the apparent FRET efficiency and the ratio CFP/YFP for the FRET between the L10-CFP **(A)** or S15-CFP **(B)** and the KCNQ2/3-YFP in the conditions with or without BACE1 or with BACE1-4C/A. **(C)** Relationship between the apparent FRET efficiency and the ratio CFP/YFP for the FRET between the L10-CFP and KCNQ2/3-YFP after treating cells with acutely-applied 5 mM MβCD, 1 mM chloroform and 0.1 mM 2-BP. The no-treatment control condition, which is the same fitted curve as in the red solid curve in panel A, is shown as the dashed curve for clarity.

Several additional pharmacological manipulations were applied to cells that showed high FRET between L10-CFP and KCNQ2/3-YFP in the presence of wild-type BACE1 (Fig. 4 C). We applied cholesterol-extracting methyl-β-cyclodextrin (MβCD) in the medium, which could efficiently decrease plasma membrane cholesterol levels and reduce the relative fraction of raft areas (Mahammad and Parmryd, 2015). MβCD application (5 mM for 5-10 mins) decreased the FRET between L10-CFP and KCNQ2/3-YFP considerably, suggesting that this FRET is dependent on the domains as well as on cholesterol that helps to maintain the rafts (Fig. 4 C). Inhaled general anesthetics also disrupt lipid rafts at anesthetic concentrations (Pavel et al., 2020). We found that one general anesthetic chloroform (1 mM) dramatically suppressed the FRET between L10-CFP and KCNQ2/3-YFP, similar to the effect of MβCD. In addition, 2-BP (0.1 mM), an inhibitor of palmitoylation, abolished the FRET, consistent with a role of palmitoylation in targeting membrane proteins to lipid microdomains.

## DISCUSSION

Using FRET, as a spectroscopic distance ruler, we found that KCNQ2/3 channels are in proximity to lipid-raft markers and to raft-specific proteins. We propose that neuronal KCNQ2/3 channels are localized to or close to the ordered membrane microdomain lipid rafts. This localization is achieved by the interaction between KCNQ2/3 and BACE1, and is dependent on the S-palmitoylation of 4 cysteines in the BACE1 carboxyl terminus. Without BACE1, KCNQ2/3 channels at the plasma membrane might be distributed more in the disordered non-raft areas at least in our heterologous expression system. Palmitoylation alters the ion channel localization and affects KCNQ channels indirectly via their auxiliary subunit BACE1 in this case. This type of ion channel regulation by S-palmitoylation is distinct from the direct lipidation and anchoring of ion-channel domains to lipid bilayers, thereby changing its gating; for instance, palmitoylation of the P2X_7_ ATP receptor prevents its desensitization (McCarthy et al., 2019). However, this indirect regulation is analogous to the inhibition of the cardiac sodium/potassium pump by palmitoylation of the auxiliary phosphoprotein phospholemman (Howie et al., 2014).

Palmitoylation manifests an important way for localizing cell signaling ranging from cellular excitability to massive endocytosis (Hilgemann et al., 2018; Linder and Deschenes, 2007). For membrane proteins, S-palmitoylation is reversible and controlled by Asp-His-His-Cys acyltransferases (e.g., plasma membrane DHHC5), thought to be abundant in ordered lipid microdomains (Howie et al., 2014; Linder and Deschenes, 2007). The palmitoylation-dependence of KCNQ2/3 localization to rafts suggests that the channel function could also be indirectly affected by this acyltransferase. Specifically, it is reasonable to predict that KCNQ channel function could be modulated by cellular signals that are upstream of DHHC, like generation of reactive oxygen species (ROS) (Gamper et al., 2006), opening of mitochondrial permeability transition pores (PTPs), coenzyme A (CoA) release, etc. (Hilgemann et al., 2018; Lin et al., 2013). Furthermore, in native neuronal environments, KCNQ2/3 channels are localized to axons by interacting with palmitoylated ankyrin G, which itself could possibly recruit KCNQ2/3 channels to lipid rafts (Chung et al., 2006; He et al., 2012). This ankyrin-G interaction could also facilitate the interaction of KCNQ2/3 with BACE1. The possible synergistic effect of the palmitoylations of KCNQ2/3-associated proteins including ankyrin G, BACE1 and AKAP79 remains elusive.

This research provides a new function for the raft-localization of BACE1, perhaps independent of the proteolytic or amyloidogenic features of BACE1. Previous research has shown that abolishing the raft-localization of BACE1 reduces the amyloid production and mitigates memory deficits in transgenic mouse models of Alzheimer’s disease (Andrew et al., 2017). Conversely, targeting BACE1 to lipid rafts by a glycosylphosphatidylinositol (GPI) anchor elevates the β-site processing of the amyloid precursor protein (Cordy et al., 2003). In addition, a recent study shows that targeted deletion of astrocyte cholesterol synthesis significantly reduces the amyloid burden in a mouse AD model, by decreasing the APP presence in lipid rafts where β- and γ-secretases are localized and functioning (Wang et al., 2021). Despite these findings, this role of the raft-localization of BACE1 for APP processing is still debated; other research suggested that the Aβ production was still present even in the absence of the palmitoylation-dependent targeting of BACE1 to lipid rafts (Vetrivel et al., 2009). Here, the raft-localization of BACE1 for the KCNQ2/3 channel function could play a role in reducing neuronal electrical excitability (Carver et al., 2020), modulating afterhyperpolarization potentials (AHP) (Gu et al., 2005; Tzingounis and Nicoll, 2008) and maintaining the proper propagation of nerve spikes (Yue and Yaari, 2004). It remains to be determined whether KCNQ2/3 channel activity affects the ability of BACE1 to cleave APP and to generate amyloid-β peptides reciprocally.

Disruption of lipid rafts is proposed to underlie the mechanism for the medical effect of inhaled general anesthetics (Pavel et al., 2020). Lipid-soluble general anesthetics change the lipid organization of sphingomyelin- and cholesterol-rich rafts and displace phospholipase D (PLD) from rafts to non-raft areas, which in turn generates phosphatidic acid (PA) that activates K_2P_ (TREK) ion channels (Pavel et al., 2020). Here, applying chloroform was able to decrease the FRET between KCNQ2/3 with the raft-marker L10, indicating that general anesthetics similarly displace KCNQ2/3 to non-raft areas and could possibly alter the activity of the channels. Future research will be needed to further understand the detailed structural mechanism and the electrophysiological effects underlying the modulation of KCNQ2/3 channels by BACE1, including whether and how signaling lipids such as PI(4,5)P_2_ and other AD-related molecular components could play regulatory roles.

## MATERIALS AND METHODS

### Cell Culture, Molecular Biology and Reagents

The tsA-201 cells were cultured, maintained and transfected as previously reported (Dai et al., 2016). The mBACE1 construct was obtained from Addgene (Cambridge, MA). The fluorescent L10 and S15 constructs were made in the lab of Dr. Byung-Chang Suh at Daegu Gyeongbuk Institute of Science and Technology (South Korea) (Myeong et al., 2021). The caveolin 1 construct was from Suzanne F. Scarlata (Worcester Polytechnic Institute, MA). Point mutations were made using Quickchange II XL Site-Directed Mutagenesis kit (Agilent technologies, Santa Clara, CA). The human KCNQ2 and human KCNQ3 constructs were fused with sequence of the enhanced YFP at the carboxyl-terminal end. The sequences of the DNA constructs were confirmed by fluorescence-based DNA sequencing (Genewiz LLC, Seattle, WA). The pANAP plasmid (Addgene, Cambridge, MA) contained the orthogonal tRNA/aminoacyl-tRNA synthetase specific to L-Anap. L-Anap (AsisChem, Waltham, MA) was dissolved in ethanol as a 10 mM stock, stored at −20°C and diluted by 500-fold into culture medium. 2-bromopalmitate (2-BP) was delivered with fatty acid-free bovine serum albumin (Sigma-Aldrich, A6003) at a stock concentration 1 mM albumin with 5 mM 2-BP, which was diluted by 50-fold into the culture medium generating a final concentration of 0.1 mM 2-BP.

### Quantification of FRET and Spectroscopic Analysis

A Nikon Eclipse TE2000-E inverted microscope with a 60X, 1.2 NA water immersion objective was used for fluorescence measurements. Epifluorescence recording was performed with wide-field excitation using a Lambda LS Xenon Arc lamp (Sutter Instruments). For spectral measurements, images were collected by a spectrograph (Model: 2150i, 300 g/mm grating, blaze = 500 nm; Acton research, Acton, MA) mounted between the output port of the microscope and an Evolve 512 EMCCD camera (Photometrics, Tucson, AZ) as previously reported (Dai et al., 2019). A filter cube containing a 376 (centered)/30 (width) nm excitation filter was used for L-Anap. CFP was excited with a filter cube containing a 425/40 nm excitation filter whereas YFP was excited with a filter cube containing a 490/10 nm excitation filter. Emission filters were removed from the filter cubes while the spectrograph functioned similarly to an emission filter to only measure the fluorescence with the light wavelength ranging from 424 nm to 583 nm (Fig. S2A). The dichroic mirror cutoffs were approximately 450 nm for the CFP filter cube, 514 nm for the YFP filter cube, and 400 nm for the Anap filter cube. CFP or YFP excitation light (not shown in figures) were also detected in the spectral images because the dichroic could not block all the excitation light, but did not interfere with the acquired fluorescence emissions of interest. Spectral images were collected with exposure times ranging from 200 to 500 ms depending on the fluorescence intensity of the individual cell using the Evolve 512 EMCCD camera and MetaMorph software (Molecular Devices, Sunnyvale, CA). The exposure time was maintained the same for recording donor emission and acceptor emission spectra of the same FRET experiment. The spectral images were analyzed with ImageJ/Fiji (National Institutes of Health). The background fluorescence spectrum was subtracted using areas outside of the cell of interest. All microscopic experiments were done at room temperature.

FRET donor-only (Anap or CFP) constructs were used to obtain the averaged donor emission spectrum excited by donor excitation wavelength. We then normalized this template emission of donor alone, called F_donor alone_ ^donor^, to the peak intensity of the F_CFP/YFP_ ^CFP^ or F_Anap/YFP_ ^Anap^ spectrum for each experiment. For the labeling used here and in other sections of the paper, the part in superscript always means the property of the excitation light, either specific to the donor or to the acceptor; whereas the part in subscript means the fluorophores expressed for the experiment, either donor/acceptor both are present, or donor alone or acceptor alone. For instance, F_CFP/YFP_ ^CFP^ indicates the emission spectrum generated after exciting the CFP when both CFP and YFP containing constructs are expressed. The normalized F_donor alone_ ^donor^ was then subtracted from either the F_CFP/YFP_ ^CFP^ or the F_Anap/YFP_ ^Anap^ spectrum in each case, generating the spectrum F_FRET_, according to the equations:

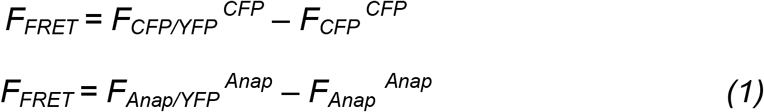

This subtraction was for correcting the bleed-through of the donor emission into the spectral range for the acceptor emission. Ratio A was then calculated as:

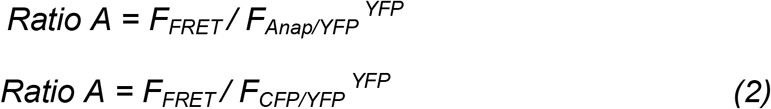

for the Anap/YFP pair or for the CFP/YFP pair. The denominator of the Ratio A is the spectrum generated by using the excitation light at the YFP excitation wavelength. In addition, two emission spectra were collected from cells that were only transfected with YFP-containing constructs (in the absence of FRET donors): F_YFP_ ^CFP^ or F_YFP_ ^Anap^, which was the emission spectrum using donor (CFP or Anap) excitation; and F_YFP_ ^YFP^, which was the emission spectrum using acceptor (YFP) excitation. Ratio A0 was calculated using:

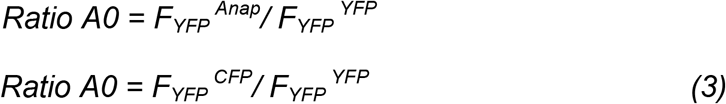

for the Anap/YFP FRET pair or for the CFP/YFP FRET pair. Ratio A0 measures the efficiency of the crosstalk of FRET, which is the fraction of the direct excitation of FRET acceptors by the donor-excitation light. The other component of such crosstalk, which is the direct excitation of FRET donors by the acceptor-excitation light is negligible for the FRET pairs used in this paper. Ratio A0 is a constant for each FRET pair; it is 0.06 for the Anap/YFP pair or 0.136 for the CFP/YFP pair. The FRET efficiency to measure was linearly proportional to the difference between Ratio A and Ratio A0; Ratio A – Ratio A0 > 0 indicated a detectable FRET signal (Erickson et al., 2001; Takanishi et al., 2006). The apparent FRET efficiency (*E*_*app*._) was calculated using the following equation 4:

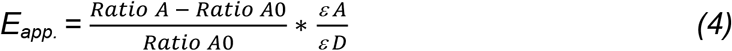

ε_A_ and ε_D_ are the molar extinction coefficients of acceptors and donors at the donor excitation. The values of ε_A_/ε_D_ were estimated as 0.10 for the CFP/YFP pair and 0.24 for the Anap/YFP pair. The properties of extinction coefficient (molar absorptivity) were from previously-measured values and from the website (www.fpbase.org): 17,500 cm^-1^ M^-1^ for L-Anap; 83,400 cm^-1^ M^-1^ for eYFP; 32,500 cm^-1^ M^-1^ for eCFP. Using the known spectrum of the YFP absorption, the ratio of YFP absorption at the donor excitation relative to the peak YFP absorption was 0.05 for the Anap/YFP pair and 0.04 for the CFP/YFP pair.

The apparent FRET efficiency *E*_*app*._ increases with the ratio of the FRET donor to acceptor peak fluorescence intensity: *F = F* _*donor*_ */ F* _*acceptor*_. At very high *F*, the *E*_*app*._ is expected to saturate and reach a steady-state value, which should correspond to the maximal FRET efficiency *E*_*max*._ (Takanishi et al., 2006). In theory, for the sensitized-emission type of FRET:

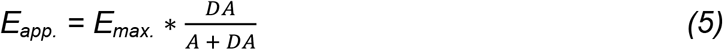

where DA and A means concentrations of the donor-acceptor pairs and the free acceptors, respectively (Takanishi et al., 2006). The quantitative amount of DA and A cannot be measured in our case. Nevertheless, in the extreme condition where all the acceptors are paired with the donors, the value A would be zero, and *E*_*app*._ equals *E*_*max*._. This equation is in contrast to the analogous equation 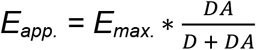 when performing FRET experiments by monitoring the donor de-quenching after acceptor photobleaching (Takanishi et al., 2006). In practice, no matter which method to perform FRET, the apparent FRET efficiency is always smaller than the real FRET efficiency *E*_*max*._.

Assuming random pairings between the donors and acceptors based on a simplified model of protein interactions (Bykova et al., 2006), and inspired by the equation 5, the apparent FRET efficiency *E*_*app*._ can also be expressed using:

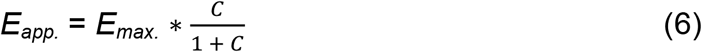

where *C* is the concentration ratio of the total donor versus the total acceptor, which is dependent on the expression level of proteins: *C = C* _*donor*_ */ C* _*acceptor*_. There is a *S* factor (*S = S* _*donor*_ */ S* _*acceptor*_) that links the molar ratio *C* to the fluorescence ratio *F* of the measured FRET donor to acceptor peak intensity: *F = C* ∗ *S*. Rearranging the equation 6 using *F = C* ∗ *S* generates the following equation for fitting the relationship between the apparent FRET efficiency and the *F*:

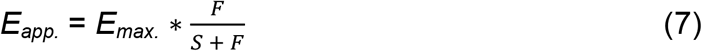

here the coefficient *S* is dependent on the optical properties of fluorophores, the recording system, the percentage of fluorophores that pair in close distance for FRET, as well as the efficiency of FRET. In general, higher *E*_*max*._ often lead to dimmer donor fluorescence and brighter acceptor fluorescence, and a resulting smaller *S*. We compared the FRET at different conditions using the *E*_*max*._ values obtained after curve fitting. The fittings were evaluated using the chi-square goodness of fit test and all the chi-square values were less than 0.1. The equation 7 is relatively model independent and serves as a simple way to predict the FRET efficiency. In addition, this equation is mathematically identical to a nonlinear saturable isotherm which was previously used for fitting the FRET data with unfixed donor/acceptor ratio (Zacharias et al., 2002): *E*_*app*._ *= E*_*max*._ *\* (F / F + K)*, where *K* is analogous to the dissociation constant for ligand binding (Kenworthy and Edidin, 1998).

The distances *r* between the donors and acceptors were estimated using the *Förster* equation: *E*_*max*._ = *1 / (1 + (r / R*_*0*_)^*6*^), and assuming the *Förster* radius R_0_, at which the FRET efficiency is 50%, to be 50 Å for the CFP/YFP FRET pair. The R_0_ value for the L-Anap/YFP pair was calculated as 47 Å using the following equation 8:

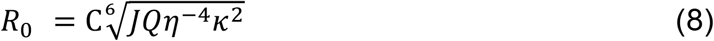

where *C* is the scaling factor, *J* is the overlap integral of the Anap emission spectrum and the YFP absorption spectrum, Q is the quantum yield of the L-Anap (approximately 0.3 for L-Anap near the cytosolic edge of the membrane) (Dai et al., 2019), *η* is the index of refraction, and *κ*^*2*^ is the orientation factor. *η* was assumed to be 1.33, and *κ*^*2*^ was assumed to be 2/3.

## ACKNOWLEDGMENTS AND FUNDING SOURCES

I thank Drs. Bertil Hille and William N. Zagotta (University of Washington School of Medicine) for support, sharing equipment, and advice on the manuscript; Dr. Donald W. Hilgemann (University of Texas Southwestern Medical Center) for helpful advice; Jongyun Myeong and other members of the Hille lab and the Zagotta lab for discussion and suggestions. This work was supported by grants from the National Institutes of Health of the United States of America: R37NS08174 and R01EY010329.

## FIGURE LEGENDS

**Figure S1.**
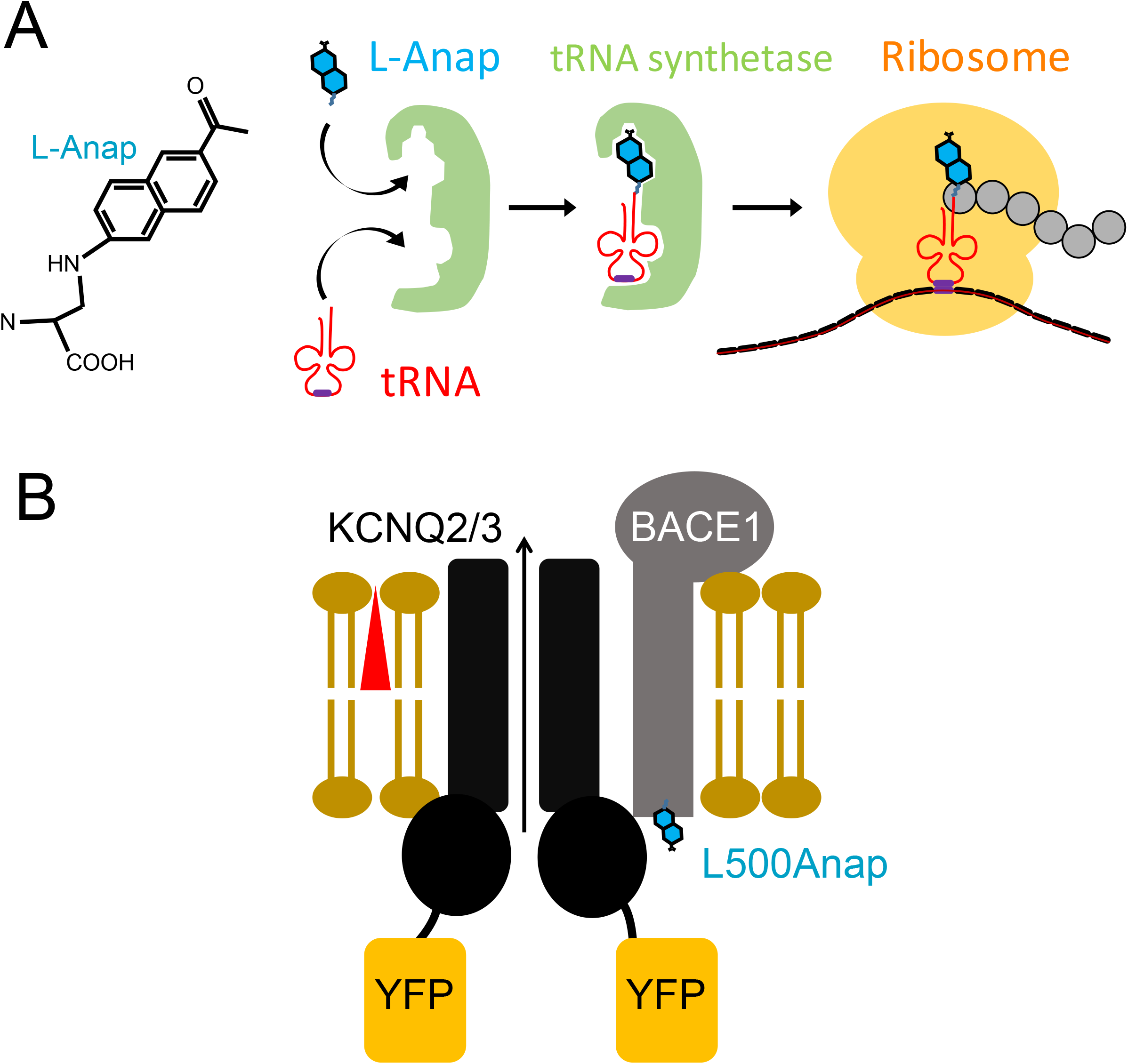
Schematic illustration of L-Anap incorporation to BACE1. **(A)** Structure of L-Anap and the Amber stop-codon suppression strategy for incorporating L-Anap. **(B)** Cartoon showing the FRET between BACE1-L500Anap and KCNQ2/3-YFP.

**Figure S2.**
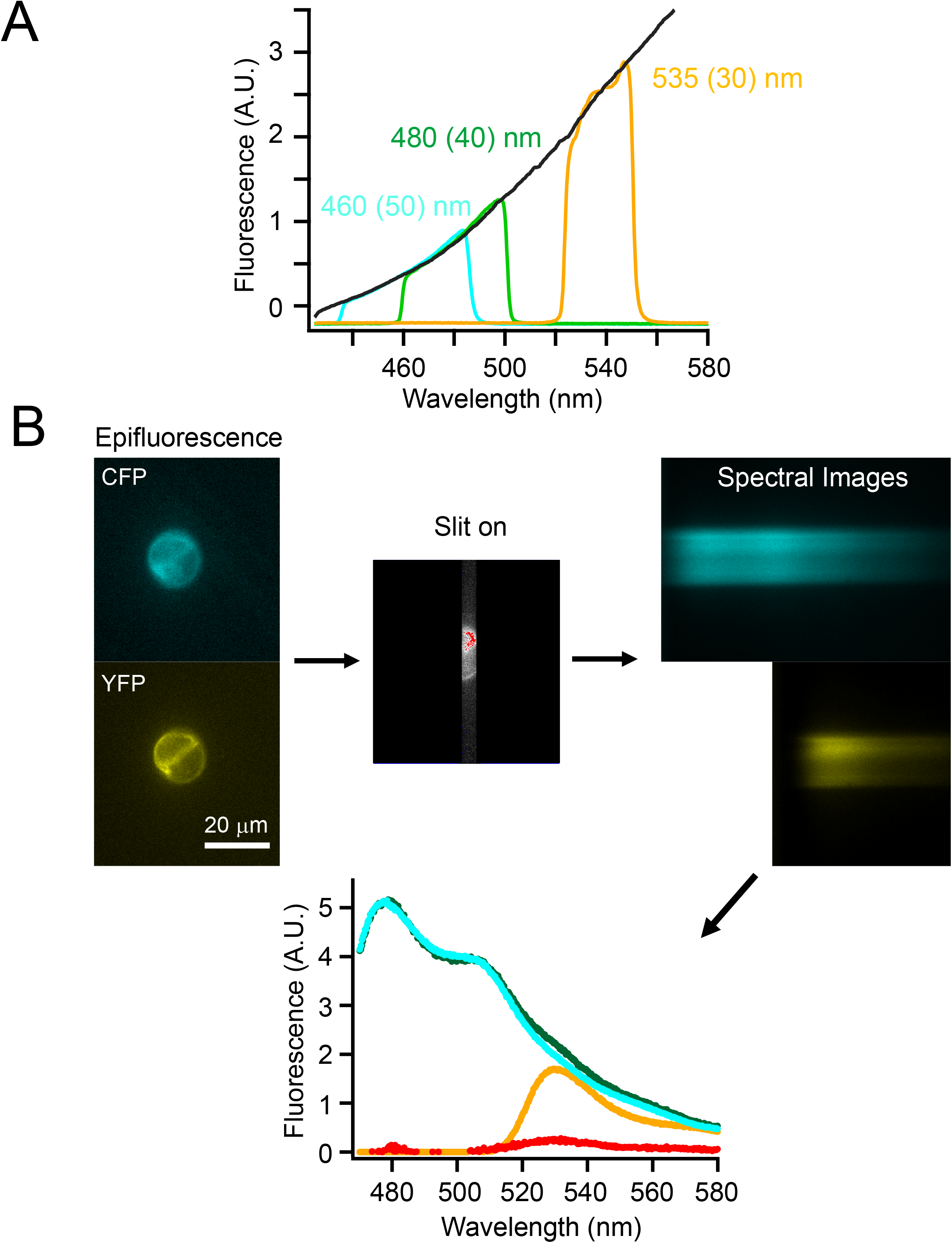
Procedure of performing spectral measurements of cultured cells. **(A)** Calibration of spectral measurements using different emission filters with Tungsten light illumination. **(B)** Flow-chart illustration of an example of measuring spectral FRET using CFP-tagged and YFP-tagged constructs transfected into tsA cells.

